# Nucleotide Asymmetry and Flexible Linker Dynamics Modulate Drug Efflux Cycle of P-glycoprotein, a Computational Study

**DOI:** 10.1101/2025.06.15.659754

**Authors:** Sungho B. Han, Jim Warwicker, Hao Fan, Stephen M. Prince

**Affiliations:** School of Biological Sciences, Faculty of Biology, Medicine and Health, The University of Manchester, Oxford Rd, Manchester M13 9PL, United Kingdom; Bioinformatics Institute (BII), Agency for Science, Technology and Research (A*STAR), Singapore, 30 Biopolis Street #07-01 Matrix Singapore 138671

## Abstract

Despite advancements in oncology, multidrug resistance (MDR) mediated by P-glycoprotein (P-gp/ABCB1) remains a major barrier to chemotherapy. P-gp is an ATP-binding cassette transporter that undergoes nucleotide-driven structural rearrangements to efflux chemotherapeutics, but the mechanistic details of the substrate transport remain poorly resolved. Here, we performed high-throughput molecular dynamics to simulate P-gp in a lipid bilayer (totaling ∼110 µs) to dissect nucleotide-dependent conformational changes across the transport cycle. Our adaptive sampling strategy reveals asymmetric nucleotide coordination at nucleotide-binding sites (NBS), which correlates with transmembrane domain (TMD) restructuring for substrate efflux. The experimentally unresolved flexible linker transiently forms up to five turn α-helix that affects nucleotide binding domain (NBD) dimerization process. We identified conformation-dependent substrate/allocrite pathways including nucleotide-specific access routes, while TMD-linker interaction facilitates substrate access tunnel formation. Together, these pathways reveal that the concerted interplay of nucleotide occupancy, linker dynamics, and overall protein conformation governs the structural plasticity and broad substrate promiscuity of the substrate binding cavity in P-gp. By integrating these findings, this work bridges static structural data with dynamic functional insights to further our understanding of the P-gp substrate translocation cycle.

## Introduction

ATP-binding cassette (ABC) transporters are integral membrane proteins that drive active transport of substrates against the thermodynamic gradient by harnessing energy from ATP molecules (1,2). This ATP-dependent transport mechanism enables ABC transporters to fulfill critical roles in diverse physiological processes, including detoxification, lipid trafficking, and immune response (3,4). In humans, overexpression of ABC transporters like P-glycoprotein (P-gp/ABCB1) compromises chemotherapeutic agents by reducing the intracellular concentration of drugs through active efflux. Understanding the structural dynamics and energetics of ABC transporters is crucial for developing adequate strategies to overcome MDR and improve therapeutic outcomes.

P-glycoprotein (P-gp) is the first drug resistance-related ABC transporter to be identified, characterized as a 170 kDa glycosylated protein capable of effluxing diverse chemotherapeutics (5). The overexpression of P-gp in tumors such as myelogenous leukemia, liver cancer and lung cancer correlates with poor chemotherapeutic treatment outcomes (6,7), as P-gp expels hydrophobic drugs (e.g., paclitaxel, doxorubicin) out to the extracellular space (8). Beyond oncology, P-gp is ubiquitously expressed at blood-tissue barriers (e.g., liver, kidney, brain) and in immune cells, where it regulates the bioavailability of analgesics, antivirals, and other therapeutics (8). Polyspecificity of P-gp to transport structurally unrelated substrates stems from a large and flexible substrate-binding cavity within the transmembrane domain (TMD). However, the lack of clear structure-activity relationships for P-gp substrates has hindered the design of selective inhibitors.

P-gp is encoded as a single polypeptide comprising two TMDs that each contains six α-helices and two nucleotide-binding domains (NBDs) (2). The TMDs form a hydrophobic cavity accessible to substrates from the intracellular milieu in the inward-facing (IF) state (9,10). During the transport cycle, the NBDs dimerize as P-gp adopts the outward-facing (OF) state, releasing the substrate into the extracellular space. Recently, a cryo-electron microscopy (cryo-EM) study revealed IF P-gp with an occluded NBD dimer interface, highlighting the conformational plasticity of P-gp undergoing IF-to-OF transition (11). The NBDs harbor conserved motifs critical for ATP binding and hydrolysis: Walker A binds a phosphate group of the bound nucleotide via a conserved lysine (GxxGxGK(S/T); x=any residue); Walker B (hhhhDE; h=hydrophobic residue) coordinates catalytic Mg²⁺ and facilitates ATP hydrolysis through a conserved aspartate; ABC signature motif (LSGGQ) from the opposing NBD stabilizes the γ-phosphate of ATP while NBDs are dimerized; adenosine-binding loop (A-loop) positions adenine ring of the nucleotide via π-stacking with a conserved tyrosine (12). ATP binding ultimately leads to NBD dimerization, sandwiching ATP molecules between the Walker A and the signature motif to enable ATP hydrolysis, which is followed by ADP and inorganic phosphate (Pi) release that resets the transporter (2,13). Several cryo-EM studies indicate that P-gp predominantly resides in IF conformations with OF states being transient (11,14,15), yet the sequence of ATP-driven transitions linking these states remains unresolved. In particular, (i) how the occupancy of the two non-equivalent nucleotide-binding sites (NBSs) alternate during the transport cycle, (ii) how the unresolved flexible linker between NBD1 and TMD2 affects conformational transitions, and (iii) the sequence of structural rearrangements linking ATP hydrolysis to substrate efflux remain poorly characterized. Combining high-throughput MD with adaptive sampling strategy, we explore the interplay between the nucleotide state and linker flexibility to gain structural and mechanistic understanding of the P-gp transport cycle.

## Results

P-gp exhibits basal ATPase activity in the absence of substrates (16), indicating that nucleotide occupancy alone can bias the transporter along its conformational cycle. To elucidate how nucleotide states modulate structural dynamics across IF and OF states, we conducted an extensive series of multiple-replica molecular dynamics (MD) simulations of human P-gp in various nucleotide states (apo, ADP/ADP, ADP/ATP, ATP/ADP, ATP/ATP) (Table 1,2). To dissect mechanistic links between nucleotide incorporation and conformational shifts in P-gp, we initiated simulations from two IF (IF-wide and IF-narrow; varying in NBD separation distance) and two OF (OF-open and OF-closed; varying in cavity access to extracellular solvent) conformations (Fig. 1a).

**Figure 1.**
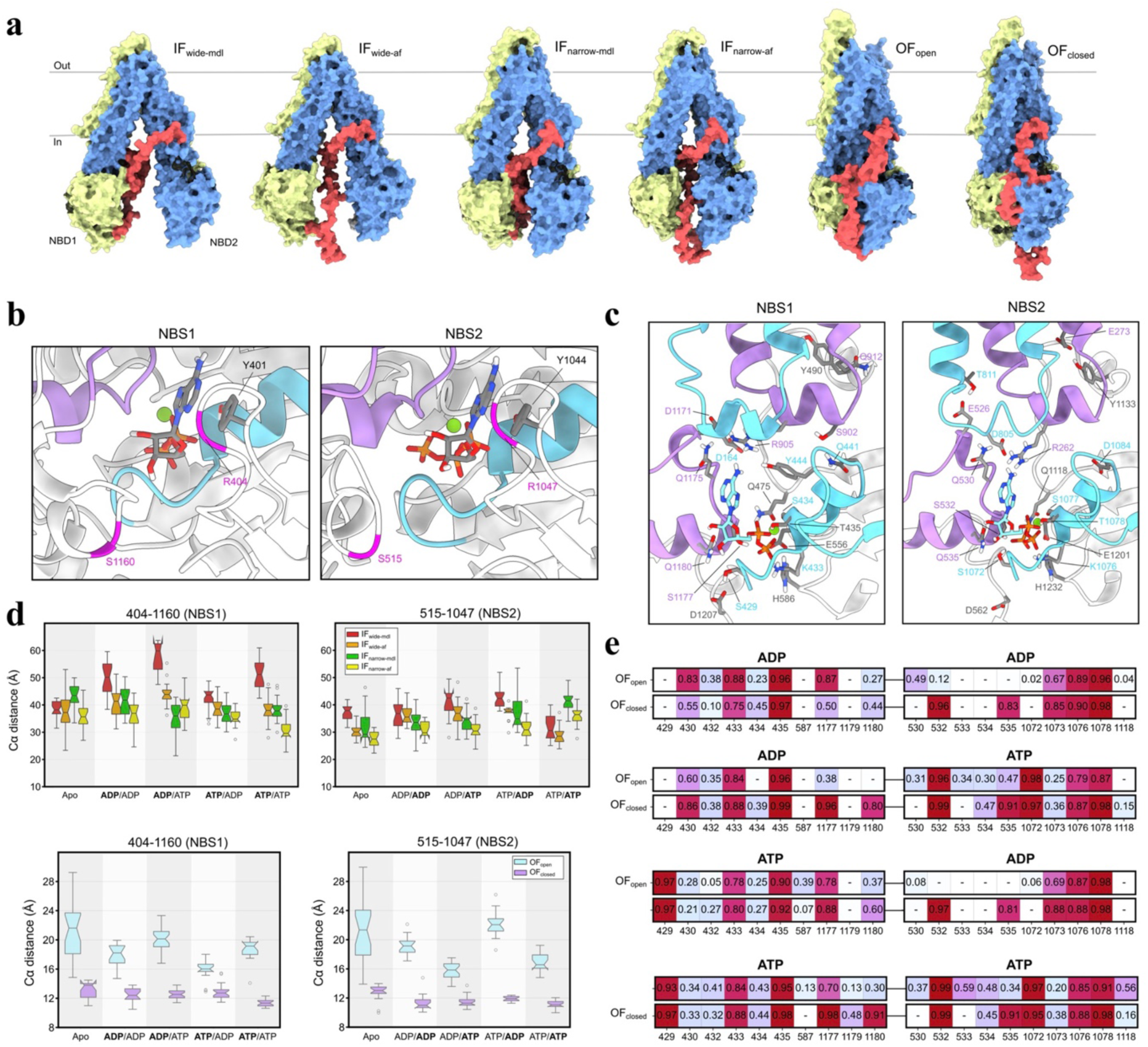
Asymmetry of the and preferential binding toward ATP and flexible linker affects nucleotide affinity at NBS. (a) Starting conformation of the four IF P-gp and two OF P-gp human homology models. The two transporter halves are colored in yellow and blue, and the flexible linker in red to highlight the differences between each P-gp starting configuration. Each of the models were incorporated with four nucleotide states; ADP/ADP, ADP/ATP, ATP/ADP, ATP/ATP. (b) Orientation of A-loop is shown, where Y401 and Y1044 are involved in coordinating the nucleotide adenosine ring. Walker A motifs (cyan), signature motif (purple) and the respective A-loop residue pairs in NBS (magenta) are shown. (c) H-bond interaction profile between the bound nucleotide and P-gp NBS. The relevant residues involved in the interaction are labelled, where the ICL1/3 and ICL2/4 are colored in cyan and purple, respectively. (d) Box plots representing the respective distributions of the residue pairs. (e) Frequency for key polar interactions between bound nucleotides and NBS residues in OF P-gp normalized over all replica trajectories and displayed as a heatmap.

**Table 1.** Total number and length of initial 10 ns multi-replica MD simulations performed in this study. All replicas were initialed with a random seed of initial velocities from the same starting configuration.

| Conformation | IF |  |  |  | OF |  |
| --- | --- | --- | --- | --- | --- | --- |
| Model PDB | wide |  | narrow |  | open | closed |
| Linker model | mdl | af | mdl | af | mdl | mdl |
| No. of nucleotide system | 5 | 5 | 5 | 5 | 5 | 5 |
| No. of replica simulation | 120 | 120 | 120 | 120 | 120 | 120 |
| Run time (ns) | 10 | 10 | 10 | 10 | 10 | 10 |
| Total run time (μs) | 6 | 6 | 6 | 6 | 6 | 6 |

**Table 2.** Total number and length of extended multi-replica MD simulations performed in this study. All replicas that displayed outlier structural deviations in any of the predefined subdomains of P-gp (see Methods) were extended to 100 ns from the initial 10 ns production run.

| Conformation | IF |  |  |  | OF |  |
| --- | --- | --- | --- | --- | --- | --- |
| Model PDB | wide |  | narrow |  | open | closed |
| Linker model | mdl | af | mdl | af | mdl | mdl |
| No. of nucleotide system | 5 | 5 | 5 | 5 | 5 | 5 |
| No. of replica simulation | 28 | 26 | 31 | 27 | 24 | 18 |
| Run time (ns) | 100 | 100 | 100 | 100 | 100 | 100 |
| Total run time (μs) | 14 | 13 | 15.5 | 13.5 | 12 | 9 |

### 1. Asymmetric Nucleotide Coordination

To probe NBS dynamics in P-gp, we surveyed residue-pair distances near the A-loop. These distances reflect conformational stability at NBS1 and NBS2 across IF and OF states under varying nucleotide conditions (Fig. 1d), as utilized in DEER spectroscopy measurements of mouse P-gp in multiple nucleotide states (17).

#### Inward-Facing Conformers Exhibit NBS1 flexibility

In IF P-gp, the A-loop residue-pair distance at NBS1 fluctuated extensively (∼25–65 Å) with maximal variability observed in the IF-wide-mdl conformer, while NBS2 exhibited comparatively restricted dynamics (∼25–45 Å) (Fig. 1d). This asymmetry suggests NBS1 adopts greater structural plasticity when NBDs are separated. The nucleotide binding energy calculations across IF conformers revealed that ATP tended to stabilize both NBSs relative to ADP; however, at NBS1 this difference was marginal except in the ATP/ATP system (Fig. S4). Protein-nucleotide electrostatic interaction analysis highlighted stable coordination of nucleotide phosphate groups by the Walker A residues S434/S1077 in all IF states (Fig. S3). Notably, ATP bound at NBS1 in the IF-narrow conformers formed an additional polar contact with T435, while purine ring of ATP interacting with R905 in intracellular coupling-loop (ICL) 4 correlated with reduced A-loop distances, suggesting nucleotide positioning modulates NBS geometry.

#### Outward-Facing Conformers Reveal Nucleotide Sensitivity in NBS2

In OF P-gp, nucleotide chemistry dictated the NBS dynamics. For “odd” nucleotide states (ADP/ATP or ATP/ADP), ATP-bound NBS exhibited closer configurations as displayed in Fig. 1d (median A-loop distances at NBS2: 15.5 Å for ATP bound versus 25 Å for ADP bound). Conversely, “even” nucleotide states (ADP/ADP or ATP/ATP) showed symmetric distances (∼18 Å) at both NBS. This asymmetry was exclusive to the OF-open conformer, with NBS2 displaying greater sensitivity between ADP and ATP in the “odd” nucleotide states (Δdistance for bound ATP and ADP: ∼10 Å in NBS2, versus Δdistance: ∼4 Å in NBS1). The observation of asymmetric nucleotide occupancy impacting NBS conformation was further corroborated by larger entropic differences between ATP and ADP in binding energies at NBS2 (Fig. S4). The OF-closed conformers lacked nucleotide-dependent A-loop distance changes, suggesting the sampled conformation is nucleotide insensitive, possibly being a post-hydrolytic state.

#### Electrostatic Interactions Underlie Nucleotide Specificity

Protein-nucleotide interaction frequency analysis revealed distinct coordination patterns for ADP and ATP (Fig. 1e). Both nucleotides formed stable interaction with Walker A residues K433/K1076 and T435/T1078 in all OF states. In contrast, γ-phosphate of ATP formed additional polar contacts with S429/S1072 (Walker A), which was absent in ADP-bound OF states. This ATP-specific interaction was replaced in ADP bound NBS with a stable interaction between β-phosphate of the nucleotide and G430/G1073 backbone. In both OF states, ATP formed more stable electrostatic interaction with NBS residues than ADP, which is in agreement with consistently lower binding energies for ATP distributing −50 kcal/mol and lower, compared to −50 kcal/mol and higher for ADP (Fig. S4). NBS2 nucleotide specificity was further evident in OF-open conformers, where the ribose hydroxyl of ATP interacted with Q535, which was absent in ADP bound NBS2. Notably, ADP bound at NBS2 in OF-open ATP/ADP state exhibited the weakest binding energy at around −110 kcal/mol (Fig. S4), suggesting ATP hydrolysis at NBS2 may initiate conformational transitions within the NBD dimer. However, the absence of significant NBD separation across OF states (Fig. S5) implies the full reversion to IF state requires extended timescales beyond the current simulations as expected.

### 2. Structural and Functional Dynamics of the Flexible Linker

The flexible linker connecting the two homologous halves of P-gp is unresolved in experimental structures due to inherent structural flexibility (9,10,18,19). To resolve the dynamic behavior of the linker, we modeled the starting configurations of the linker using two distinct approaches: (1) ab initio prediction (MODELLER, referred as “mdl”) and (2) AlphaFold2-derived secondary structure (referred as “af”) (Fig. S1).

#### Linker Architecture and Helix-Dependent Interactions

In the simulations, both models revealed that the residues 669–690 of the linker form a transient α-helical structure (Fig. 2c/d). In IF-af conformers, the helical propensity of the residues 669–690 was higher than IF-mdl conformers, likely due to the initial helical content in the starting configuration in IF-af models. Notably, we observed the linker residues 669–690 forming electrostatic interactions with TM helices 3, 9, and 10 (Fig. S6). Positively charged arginines in the linker (R669, R673, R680) formed salt bridges with the acidic TM3/10 residues, while negatively charged aspartates/glutamates in the linker (D679, E686, E689) interacted with basic TM9 residue side chains, adhering the linker in between opposing TMDs. This transient helix formation correlated with shorter ICL distances in IF-af conformers (TM3–4: ∼25–30 Å; TM9–10: ∼25– 27 Å) compared to IF-mdl conformers (TM3–4: 27–37 Å; TM9–10: 28–35 Å) (Fig. 2b).

**Figure 2.**
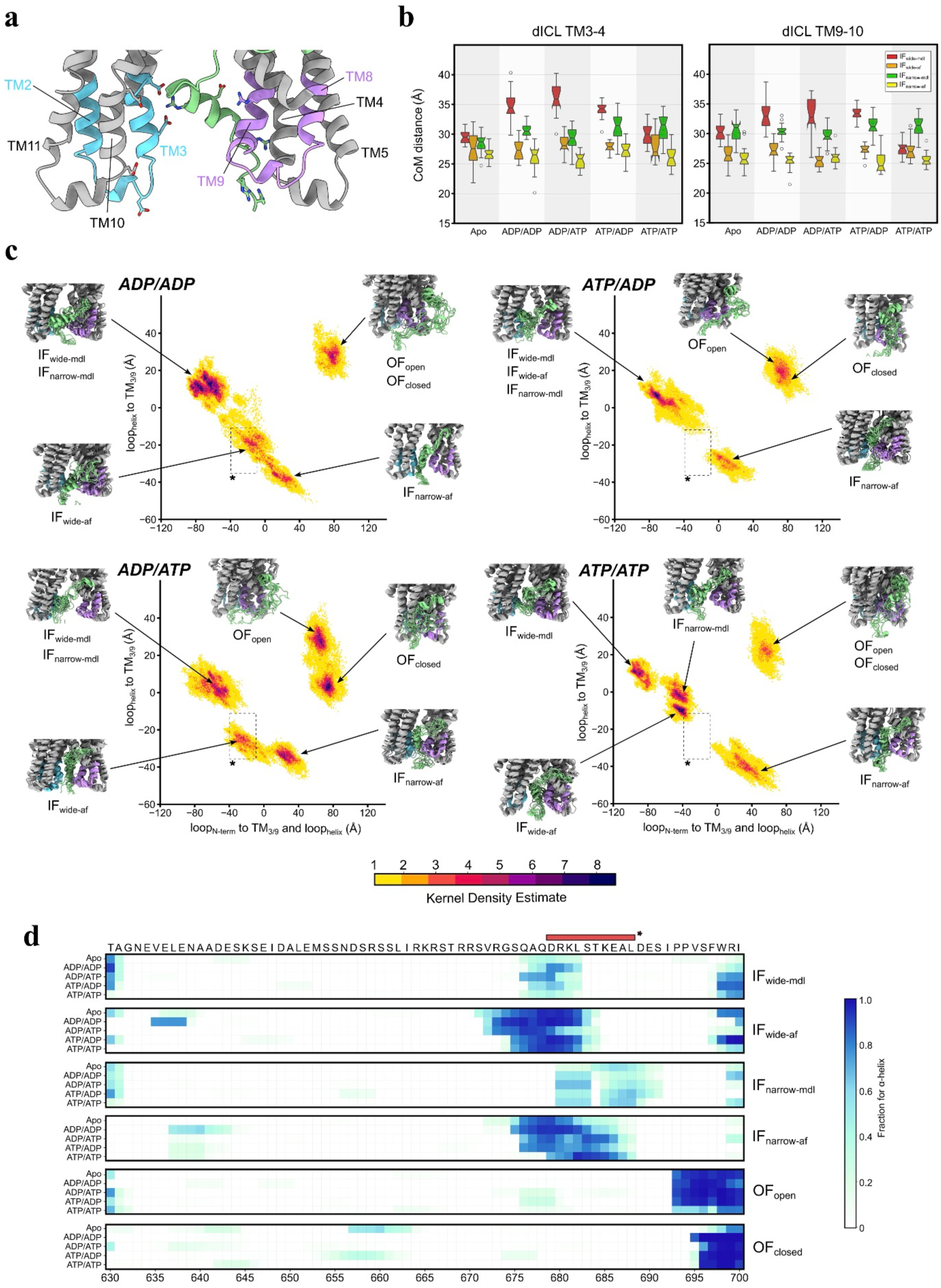
Flexible linker forms a helix and forms stable interaction with TM3/9. (a) ICLs and intracellular portions of TM helices. ICL1 and ICL4 (cyan and purple) interact with the helical segment of the flexible linker via H-bonding interactions between polar side chains. (b) Center-of-mass distance between intracellular portions of TM3-4 and TM9-10 representing the distance between ICL1-4 and ICL2-3, respectively. (c) PCA heatmap with PC1/2 for four nucleotides state. PC1 represents the distance between the N-terminal side of the linker and the ICLs and linker helix. PC2 represents the distance between the ICLs and linker helix. A region between −40 to 0 PC1 and −10 to −40 PC2 is marked with a box to highlight the difference across all PCA results. The representative structures of the linker helix-ICL region are shown for each of the six P-gp conformers in each nucleotide state. (d) Alpha helical propensity of the flexible linker (630–700) normalized over all replica trajectories shown as helicity heatmap for all six P-gp conformers in apo and four different nucleotide states. Red bar with asterisks represents the helical region (679-688) within the AlphaFold2 P-gp model, which has been the template for the linker configuration in IF-wide-af and IF-narrow-af homology models.

#### Conformational Landscape of Linker-TMD Interface

Principal component analysis (PCA) of linker-TMD conformations, visualized via PyEMMA-derived kernel density estimation (20), revealed distinct nucleotide- and conformational-dependent populations (Fig. 2c). IF-mdl conformers, characterized by lower helical content (∼10-15 %) within the linker, occupied a broad conformational space in the PCA map (top-left quadrant), representing the disordered linker configuration. In contrast, IF-af conformers were closely distributed at the bottom center region of the heatmap, representing the microstate configuration with closer TMD-linker proximity stabilized by electrostatic interactions. Interestingly, IF-wide-af conformers with NBS1 bound to ADP (ADP/ADP, ADP/ATP) resulted in the linker helix formation closer to the N-terminus (Fig. 2d), which was represented as a unique microstate density in the PCA heatmap (Fig. 2c, annotated box). These observations suggest the global P-gp conformation and possibly nucleotide state can affect conformational variability in the linker.

### 3. Conformation-Dependent Substrate Access Pathways

Having established NBS asymmetry and linker-mediated NBD/TMD coupling, we next explored how these conformations shape drug-translocation tunnels connecting the substrate binding cavity and surfaces of the membrane leaflets and the solvent.

#### IF State: Substrate Accessible Routes

CAVER analysis (21) of IF conformers revealed three ligand-accessible pathways connecting the TMD substrate-binding cavity to the intracellular lipid/solvent interface (Fig. 3): (i) tunnels between TM4 and TM6 (channels 2a, 2b) in the N-terminal half of P-gp, (ii) a lateral portal via TM10/12 (channel 2c) in the C-terminal half, and (iii) cytosolic-facing portals near the solvent interface (channels 1b, 1c). In the IF-wide-mdl conformer, the nucleotide state altered the ligand-accessible pathway availability and the separation distance between the two NBDs. Channel 2c appeared exclusively when NBS1 was bound to ADP (Fig. 3c), which correlated with higher NBD center-of-mass (CoM) distances above 50 Å (Sup. Fig. 5). Channel 2a was observed in conformers with ADP occupancy at NBS2, which had lower NBD separation distance below 50 Å. Channel 2b remained frequently open (15∼44 % of frames), supporting experimental observations that TM4/6 form a common entry portal (10,11). In contrast, IF-wide-af formed channels 2a/2b only, independent of nucleotide state (Fig. 3d). The absence of TM10/12-associated tunnels (channel 2c) and consistent occurrence of channels 2a/2b toward the intracellular solvent and lower lipid leaflet interfaces suggest linker-mediated stabilization of binding cavity access via TM4/6. IF-narrow conformers lacked detectable tunnels with adequate frequency (all occurrences were < 10% of frames) with overall lower NBD CoM distances than IF-wide (Sup. Fig. 5). This suggests the conformational microstate observed in IF-narrow conformers represents an IF-occluded state with inaccessible binding cavity and closer NBDs.

**Figure 3.**
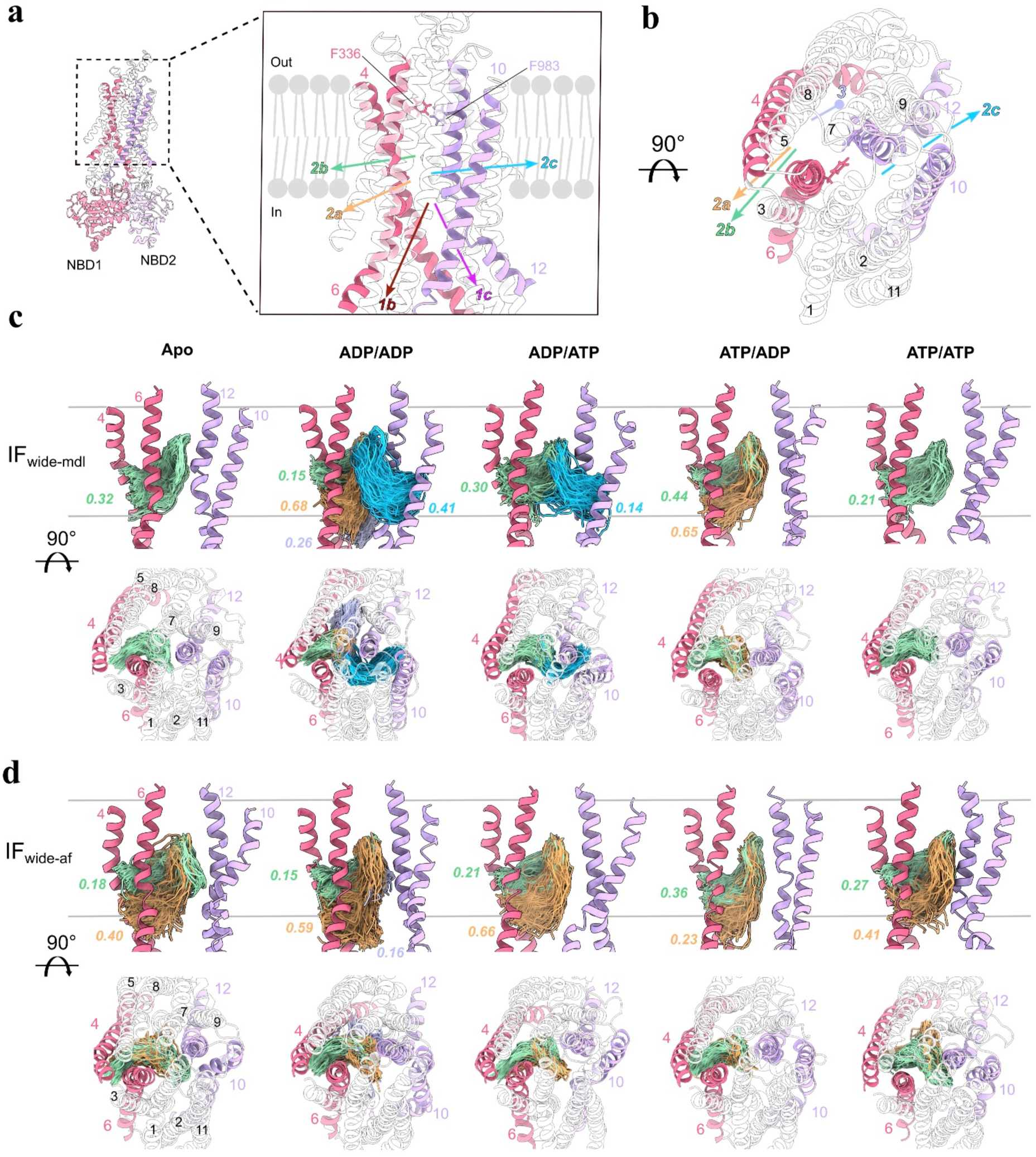
Ligand accessible tunnels are shaped in a nucleotide dependent manner in IF P-gp for potential substrate ingress into the binding cavity. (a) View from the lipid bilayer cross section highlighting the ligand access tunnels identified by CAVER 3.0. Starting position of probes were residues F336 and F983 within the cavity. Tunnels are defined as the following: 1b-intracellular opening near TM4/6 (pink); 1c-intracellular opening near TM10/12 (purple); 2a-intracellular side bordering lower leaflet access route via TM4/6; 2b-lower leaflet access route via TM4/6; 2c-lower leaflet access route via TM10/12; 3-inner sub-pocket near TM4/8. (b) Alternative view of the ligand access tunnels. (c) Nucleotide dependent ligand access gate formation in IF-wide-mdl. Along the grey lines representing lipid bilayer boundaries to solvent, relevant tunnel paths from all simulation replicas and corresponding tunnel occupancy are shown. Each occupancy value is normalized based on the total number of frames from the respective replica trajectory pool. TM4/6 (pink) forms routes to tunnels 2a and 2b and TM10/12 (purple) routes to tunnel 2c. Last 70 percent of frames were used to analyze converged portions of the trajectories. Tunnels with occupancy values higher than 0.10 are depicted. (d) Nucleotide dependent ligand access gate formation in IF-wide-af. No tunnel above 0.10 occupancy were detected in IF-narrow P-gp conformers.

#### OF State: Substrate Egress Routes

In OF conformers, three ligand tunnels were identified (Fig. 4): (i) TM1/3-bounded channels (5a, 5b), (ii) TM7/9-associated routes (4a, 4b), and (iii) a solvent-exposed portal above the binding pocket (channel 6). OF-open exhibited nucleotide-dependent formations of the substrate egress tunnels. The apo state showed limited occurrence of channel 5a (Fig. 4c), while ADP/ADP state showed tunnels toward the upper lipid leaflet and extracellular solvent interfaces via TM7/9 (channels 4a/4b) and the extracellular portal (channel 6). Notably, the ADP/ATP state produced the largest ensemble fraction of open egress tunnels in high frequency. The channels 5a/5b toward TM1/3 and channels 4a/4b toward TM7/9 appeared in ∼80% of the frames, while the solvent-exposed channel 6 was detectable in 86% of the frames. The extensive opening of the binding cavity in the ADP/ATP state correlated with significant rearrangement in the TMD, as shown by the elevated distances between TM3–11 (33.5 Å), TM6–12 (23 Å), and TM8–9 (31 Å) (Fig. 4e). This correlation hints that loss of the γ-phosphate at NBS1 may relax TMD packing, potentially lowering the barrier to substrate release. Conversely, OF-closed conformers, marked by tightly packed extracellular TM helices (e.g. TM5–11: 32 Å in ADP/ADP state), lacked detectable tunnels with adequate frequency (above 10% of the frames), which suggests OF-closed microstate being a post-transport configuration (Fig. 4e).

**Figure 4.**
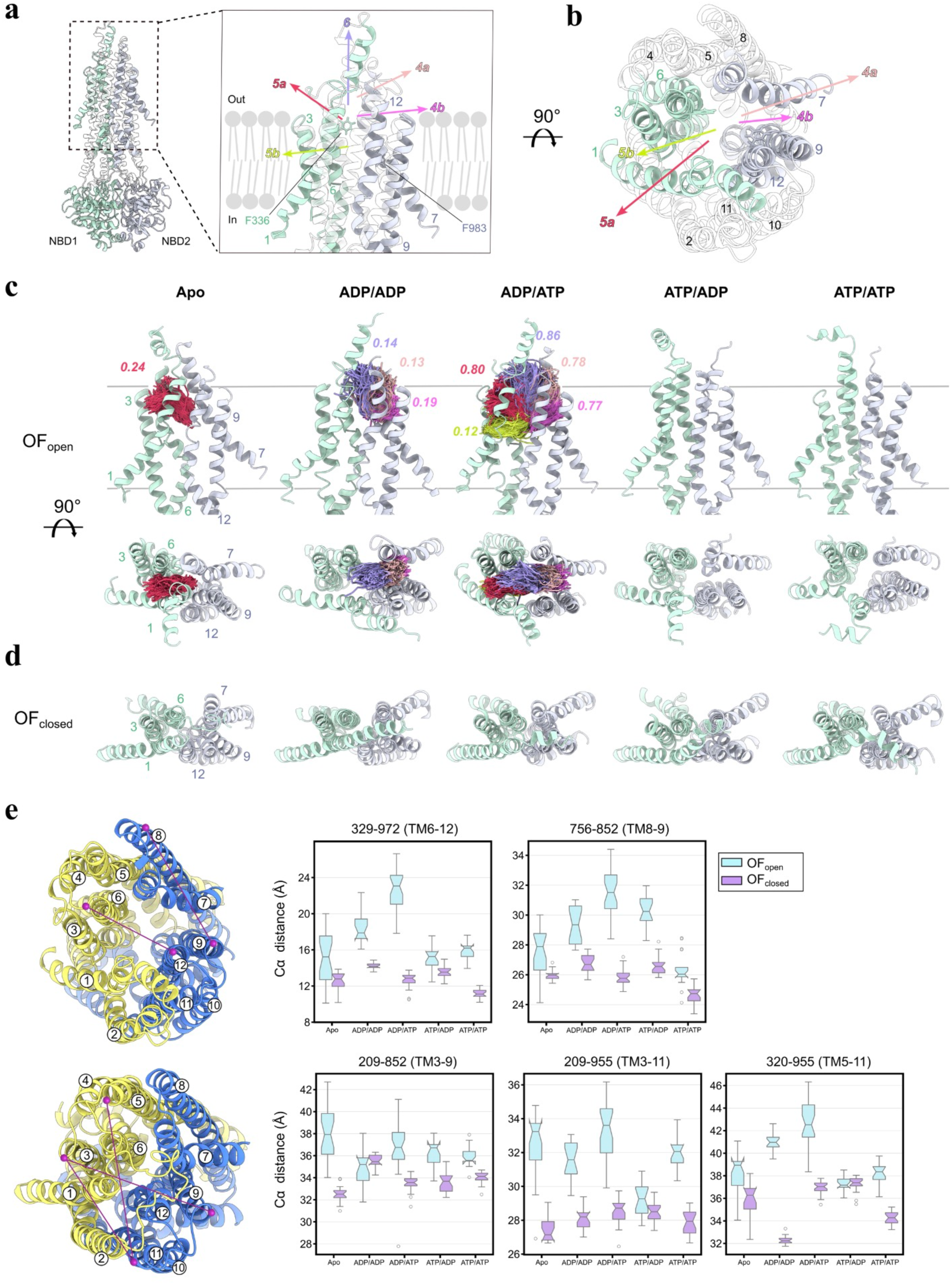
Potential substrate egress channels identified in OF P-gp showing high structural plasticity upon nucleotide incorporation. (a) Viewed from the lipid bilayer cross section as Fig. 3. Tunnels are defined as the following: 5a- extracellular side bordering upper leaflet access route via TM1/3 (green); 5b- upper leaflet access route via TM1/3 (green); 4a- extracellular side bordering upper leaflet access route via TM7/9 (blue); 4b- upper leaflet access route via TM7/9 (blue); 6- extracellular space egress toward the solvent near TM6/12. (b) Alternative view of the ligand access tunnels. (c) Nucleotide dependent ligand access gate formation in OF-open P-gp configurations. Gray lines represent lipid bilayer boundaries to intra- and extracellular space. Relevant tunnel paths from all simulation replicas and corresponding tunnel occupancy are shown. TM1/3 (green) forms routes to tunnels 5a and 5b and TM7/9 (blue) routes to tunnel 4a and 4b. Last 70 percent of frames were used to analyze converged portions of the trajectories. Tunnels with occupancy values higher than 0.10 are depicted. (d) Top view of the OF-occluded P-gp configurations. No tunnels above 0.10 occupancy were detected. (e) Extracellular residue pair distances. The two TMD halves, TMD1 and TMD2, are colored in yellow and blue, respectively. The residue pairs used in the box plots are depicted with connected dots in purple.

## Discussion

### 1. High-Throughput MD Sampling Framework

Recent advances in structural biology, including cryo-EM and deep learning-based structure prediction tools like AlphaFold, have dramatically expanded the availability of static snapshots of ABC transporters (22). However, capturing transient intermediate conformational states essential for understanding functional dynamics can be guided using computational approaches such as MD simulations, which could bridge temporal and spatial resolution gaps between experimental techniques. Here, we implement an adaptive MD workflow combining 120 short, parallel simulations followed by selective extension of replicas exhibiting outlier structural deviations (Tables 1, 2). This strategy achieves markedly broader conformational sampling compared to a single extended 1.8 µs MD trajectory (Fig. S2). Limited conformational transitions in the conventional MD (cMD) benchmark suggests kinetic trapping of P-gp configuration in the initial metastable state due to the rugged free-energy landscape, as evidenced in other MD benchmark studies (23,24). By prioritizing trajectory extension based on structural variance, similar to machine learning-guided latent space sampling approach (25), our methodology displays greater sampling of transient conformations compared to the cMD benchmark. The cMD simulations of membrane proteins like P-gp often suffer from insufficient sampling due to the high conformational degrees-of-freedom as evidenced by poor structural convergence in 3 x 200 ns simulation of mouse P-gp (26) and persistent sampling deficits of membrane bound P-gp even after 500 ns (27). Our strategy partially mitigates the stochastic noise through redundancy with the use of a high number of replica simulations and tailors simulations to explore transient states via targeted extensions of structurally more variable trajectories. This high replica MD protocol could be utilized to investigate different metastable states of other ABC transporters and other membrane proteins.

### 2. NBS Asymmetry and Mechanochemical Coupling

The asymmetric nucleotide coordination of NBS1 and NBS2, evidenced by divergent A-loop distances (Fig. 1d) and binding energies (Fig. S4), is consistent with the two-stroke (“power stroke”) model of ABC transporters, where ATP hydrolysis at one NBS drives conformational transitions (28,29). This asymmetry is further corroborated by DEER spectroscopy studies (17), which revealed two distinct populations (45 Å and 60 Å) for distances between spin-labeled residues at NBS1 in mouse P-gp, compared to a single peak (∼45 Å) at NBS2.

These experimental observations align with our simulations: In IF conformers, NBS1 exhibits higher flexibility (A-loop distance: 25–65 Å) compared to NBS2 (25–45 Å) (Fig. 1d). In OF conformers, heightened sensitivity to nucleotide anhydride chemistry in NBS2 emerges as ATP and ADP coordinates with uniquely different residues (Fig. 1e). For instance, the dimerized NBDs enable full coordination of γ-phosphate of ATP by the Signature motif (Q535) and Walker A (S1072) residues, while these interactions are absent in ADP bound NBS2. This NBS asymmetry in OF P-gp, absent in IF states, may contribute to the mechanochemical coupling between ATP hydrolysis and TMD restructuring during substrate efflux that is unique to OF conformations.

The functional divergence between NBS1 and NBS2 is underscored by mutational studies. For instance, simultaneous mutation of Walker A lysines (K433/K1076) abolishes P-gp activity, while single mutants retain partial ATPase and transport function (30). These findings align with our observation that K433/K1076 forms a stable electrostatic interaction with γ-phosphate of ATP (Fig. S3), which is absent in ADP bound states. The additional electrostatic interactions within NBS established in the presence of ATP is likely related to the thermodynamic preference for ATP over ADP. This idea is supported by free energy calculations of P-gp NBD dimers, where ATP bound states exhibit lower potential of mean force profiles than ADP-Pi bound states (31).

The NBS asymmetry in ATP hydrolysis is evidenced by biochemical studies of Walker B human P-gp mutants (D555N/D1200N) (32). Using vanadate-trapping combined with [α-32P]-8-azido-ATP photolabeling, the authors showed that under turnover conditions, the C-terminal half of P-gp (NBS2) was preferably driven into the post-hydrolysis state unlike the N-terminal site (NBS1) remaining predominantly in an ATP-bound state, suggesting ATP is hydrolyzed preferably at NBS2 than NBS1. This parallels our observation of nucleotide-specific coordination changes for NBS2 in OF-open conformers. While ATP binding stabilizes NBS2 via γ-phosphate coordination (Fig. 1e), ADP occupancy correlates with elevated A-loop distances (25 Å and 15.5 Å for ADP and ATP, respectively). The asymmetry in NBS is critical for the “alternating access” mechanism (33), where iterative ATP hydrolysis at one NBS drives TMD rearrangements. However, the absence of significant NBD separation in OF states (Fig. S5) suggests the full IF→OF→IF transitions require prolonged timescales or substrate binding. This is consistent with MD studies of Sav1866 showing stable NBD dimerization in ATP bound states but progressive dissociation in ADP bound systems (34,35).

Taken together with earlier spectroscopy and mutagenesis studies, our simulations add a concrete detail: in the inward-facing ensemble the more flexible NBS1 favours possible nucleotide exchange, whereas full γ-phosphate coordination at NBS2 emerges only after the transporter samples the outward-facing state. This site-specific difference in ATP versus ADP handling was not captured in the published experiments and refines current descriptions of the alternating-access cycle.

### 3. The Flexible Linker as a Potential Conformational Gatekeeper

Our findings position the unresolved flexible linker as an important structural component of the P-gp conformational cycle. The partially helical linker in IF-af conformers stabilizes the intracellular interface between opposing TMDs (Fig. 2d) via electrostatic networks. The salt bridge formation between the charged linker residues (R669/D679) and TM3/9/10 (Fig. S6) effectively acts as a “molecular glue”, modulating the NBD-NBD distance (Fig. S5) and TMD flexibility as evidenced by linker conformer dependent substrate tunnel formations (Fig. 3c/d). Consistent with this role, cryo-EM studies have shown that drug binding within the central cavity induces a pronounced shift of TM9, described as an “initiator of the peristaltic extrusion” mechanism (36). Our simulations further indicate that helix formation within the linker correlates with reduced inter-ICL distances (Fig. 2b). The stable interaction between the linker and TMDs could impede premature NBD dimerization and prolong substrate-binding cavity accessibility, forming an IF occluded microstate. The IF occluded-state with the collapsed binding cavity and close NBD-NBD distance has been previously observed in a bacterial exporter McjD and mouse P-gp (11,37). Interpretation of the variable helicity in the linker must, however, account for potential bias from the initial model configurations deployed in this study. The linker-mdl construct was introduced without a helical portion in the linker, whereas the linker-af model contained a pre-formed α-helix (residues 679–688) predicted by AlphaFold2 (Fig. S1). Despite these differences in the initial configuration, emergent helicity developed in IF-mdl simulations, and IF-af simulations exhibited dynamic helicity across a broader segment (residues 671–687). Notably, IF-wide-mdl trajectories displayed spontaneous helix formation in residues 673–683, mirroring the AlphaFold2-predicted region (Fig. 2d). IF-narrow-af conformers showed positional variability of the helix, despite the intrinsic α-helix-stabilizing propensity of the AMBER force field (38). In IF-wide-af conformers, the helical segment shifted toward the N-terminus in ADP/ADP and ADP/ATP states, showing that the nucleotide state can transiently affect the linker conformation. Collectively, these observations indicate that linker helicity is not a modeling artifact, but rather a dynamic feature that reorients in response to the global conformational landscape of P-gp and potentially affected by the nucleotide state.

The regulatory role of the linker has been hypothesized due to the presence of conserved charged residues in this region across ABC transporters (39). Previous biochemical studies postulated a potential regulatory role of the linker via phosphorylation (40–42), although the results are contrasting (43). Phosphorylation of serine residues (S661, S667, S671, S675, S683) within the helical linker region (residues 671–691) of P-gp could modulate its helicity, potentially fine-tuning transporter activity (41,44). Interestingly, alanine mutations of these serines do not abolish transport (41), suggesting the change in electrostatic profile of the linker does not critically affect the dimerization kinetics. Functional studies of linker truncations further support the structural role of the linker. Removal of residues 653–686 (Δ34) abolished ATPase activity in human P-gp, while reintroducing a 17 residues long glycine/serine-rich peptide into the Δ34 shortened linker partially restored function (40). This implies that the linker plays a critical role for the structural stability and conformational dynamics of P-gp. From our simulation data, we observe transient linker helicity (residues 673–690) in IF conformers (Fig. 2d) and frequent salt-bridge formation between the charged linker residues in the helical portion and the intracellular segments of TMDs (Fig. S6). We predict that the charged residues in the native linker might introduce a kinetic barrier for NBD dimerization via electrostatic tethering, affecting the conformational transition kinetics. MD simulations of murine P-gp corroborate this idea, showing the linker dampens NBD fluctuations and solvent accessibility (45,46). However, prior studies failed to observe persistent helical content in the linker due to limited sampling (45,46). Despite our efforts to enhance conformational sampling with the use of a multi-replica approach, intrinsic flexibility of the linker and the lack of inherent secondary structure mean our simulations do not exhaustively cover the full dynamic landscape of the linker, as evidenced by PCA heatmaps (Fig. 2c). Nevertheless, the consistent emergence of linker helicity across models and nucleotide states underscores the role of the linker as a potential conformational buffer, transiently modulating TMD/NBD coupling via electrostatic tethering, when localized in between the two wings of P-gp. With this insight, stabilizing the transient helical linker-TM3/9 interaction via allosteric modulator could prolong the IF-occluded intermediate state, which may slow transport activity and enhance chemotherapeutic retention.

If stabilization of the linker helix indeed steadies the inward-facing state, progression through the transport cycle would require this segment to relax or unwind so that the NBDs can finally dimerize. Such short-lived structural shifts lie outside the reach of current structural methods, thus future computational studies can shed light on this potential linker maneuver and point to experiments that test their role.

### 4. Dynamic Substrate Entry and Release Pathways

In IF P-gp, we identify formations of ligand accessible tunnels involving TM4/6 and TM10/12 (Fig. 3c), showing the flexible nature of the substrate binding pocket in P-gp to accommodate diverse substrates. The involvement of TM4 and TM10 in the formation of ligand access tunnels has been observed in cryo-EM studies of human P-gp bound to substrate/inhibitors, which showed high structural plasticity in TM4/10 that are actively engaged in trapping the drug in the binding cavity. Our simulations add an additional dimension to the ligand binding dynamics, as we observe the partially helical linker conformation stabilizing ligand accessibility to the central cavity via TM4/6 in IF-wide-af conformers (Fig. 3d), suggesting a possible facilitatory mechanism of the linker to aid substrate entry during nucleotide exchange. Although, the linker-TMD interaction and the respective binding cavity rearrangement are likely transient the occlusion of binding cavity near TM4/6 with “molecular plugs” like zosuquidar and elacridar shows high efficacy in P-gp inhibition (47–49). Nucleotide-dependent TM10/12 tunnel formation in IF-wide-mdl underscores the dynamic nature of substrate recruitment in the presence of flexible linker configurations, as ATP or ADP occupancy of NBS correlates to differential occurrence of ligand accessible channels from the central cavity.

In OF states, asymmetric nucleotide occupancy in OF-open bound to ADP/ATP shows the highest ensemble population of open efflux channels, with solvent-exposed portal (channel 6) and tunnels toward upper lipid leaflet and solvent interfaces (channels 4a, 4b, 5a) observed in most of the frames (Fig. 4c). ADP/ADP state exhibits transient cavity opening toward the extracellular solvent (channel 6: 14% occupancy) and upper leaflet via TM7/9 (channels 4a,4b: 13–19% occupancy), while the TM1/3 side of the cavity remains inaccessible. The occurrence of substrate egress tunnels in partially or fully hydrolyzed nucleotide states (ADP/ATP and ADP/ADP states, respectively) in our simulations aligns with the “ATP switch” model (50), where ATP hydrolysis at NBS drives conformational resetting. ATP/ATP and ATP/ADP states in OF-open results in no detectable egress tunnels, as the distance between the extracellular ends of TM6 and TM12 is lower (TM6–12: ∼16 Å) in the respective nucleotides (Fig. 4e) leading to tighter packing of TM helices and binding cavity. These findings align with MD studies of Sav1866, where ATP/ATP states collapsed the substrate cavity while asymmetric nucleotide states led to opening of the extracellular gate (51,52). Unlike the OF-open ensembles, our OF-closed trajectories show no detectable efflux tunnels and retain a tightly packed TM6–TM12 interface (Fig. 4e). The OF-closed model was initiated from the ATP-bound E556Q/E1201Q (“EQ”) mutant, a construct that cryo-EM and biochemical probes have assigned to an outward-occluded, post-hydrolytic state (19,53). The persistence of the sealed cavity and dimerised NBDs in our simulations, independent of the nucleotide states, supports the interpretation that our OF-closed conformer represents a post-transport intermediate in which the export pathway has already shut, and the protein awaits nucleotide release to reset.

The asymmetric role of NBS1 and NBS2 as observed in tunnel formation from the OF-open conformers is supported by biochemical studies. Hrycyna and colleagues proposed that each hydrolysis event serves distinct functions (32). ³¹P SSNMR on BmrA trapped in various nucleotide states showed one NBD occupies a high-energy post-hydrolysis conformation, while the other NBD holds a non-hydrolyzed ATP, directly reporting on asymmetric catalytic cycling (54). H/D exchange data by Vigano et al. (55) showed when E552Q and E1197Q mouse P-gp mutants are trapped in the post-hydrolysis state with ADP-Vi then rescued by adding ATP, the H/D exchange rate was increased from ∼45 to ∼80% for E552Q P-gp, but no significant change was observed in E1197Q mutant (55). With this data, the authors suggested hydrolysis at NBS1 drives major membrane-domain rearrangements. Based on the observation of substrate egress tunnels in OF-open conformers, we propose that ATP hydrolysis at NBS1 initiates substrate extrusion, while NBS2 hydrolysis resets the transporter.

## Conclusions

This study presents a high-throughput MD framework that efficiently characterizes metastable states in P-gp, partially overcoming sampling limitations of conventional simulations. By deploying parallel short replicas and selectively extending trajectories with high structural variance, we capture transient intermediates critical to substrate transport cycle of P-gp. Our simulations show a stage-wise model in which (i) NBS asymmetry biases inward facing ensembles, (ii) transient linker helicity modulates the inward facing to occluded state transition, and (iii) combined effects reshape substrate tunnels predicted for ligand entry and release. Together, our simulations link nucleotide chemistry, linker flexibility and substrate pathway to provide more detailed picture of how P-gp moves its cargo across the membrane. Beyond P-gp, our workflow provides a paradigm for probing metastable dynamics in other ABC transporters and flexible membrane proteins in general, bridging gaps between experimental insights and functional mechanism elucidation.

## Materials and Methods

### 1. Human P-glycoprotein 3D Structural Modelling

Crystal structures of IF murine and mouse P-gp (PDB: 4Q9H, 4M1M respectively) (9,10), OF Sav1866 (PDB: 2HYD) (18) and cryo-EM structure of OF human P-gp (PDB: 6C0V) (19) were selected as templates for building full length human P-gp homology models. All sequences in template structures were mapped to human P-gp sequence (Uniprot ID: P08183) and missing residues were modelled using Modeller v9.23 (56). For the IF models, a total of 5000 models of flexible linker (residues 630-699) were generated and a conformation with the highest Discrete Optimized Protein Energy (DOPE) was selected as starting configuration (referred as linker-mdl, see Fig. S1). The second starting configuration of the flexible linker was constructed by adopting the AlphaFold2 predicted structure (AF-P08183-F1-v4) (22). The linker structure for OF P-gp homology models were generated, as described for linker-mdl.

### 2. MD Simulation Set up

Protein was embedded in 5:1 1-palmitoyl-2-oleoyl-sn-glycero-3-phosphocholine (POPC):cholesterol bilayer using CHARMM-GUI (57) based on the protein-membrane orientation predicted by OPM database (58). ATP-Mg2+ were docked by aligning nucleotide bound structure of ABCB10 (PDB: 4AYT) (59). Simulations were performed using AMBER ff14SB force field for protein (60) and LIPID14 FF (61) for POPC-cholesterol membrane, and phosphate and magnesium parameters from Meagher et al. (58) and Allnér et al. (63). The periodic TIP3P water box was used with Na+ and Cl-ion concentrations of 150 mM. System was energy minimized using AMBER20 (64), applying harmonic restraints with a force constant of 1000 to 0 kcal/mol Å2 on the heavy atoms of protein, as described in other membrane simulation protocols (65). GROMACS (66) 2021.5 package was used for equilibration and production runs. NPT ensemble was used during equilibration with semi-isotropic coupling at the constant pressure of 1 atm and constant temperature of 310 K, using Nosé-Hoover Langevin piston and Langevin thermostat. Time step of 1 fs was used for the initial 2.5 ns equilibration run with harmonic restraints from 100 to 0 kcal/mol Å2, followed by 12.5 ns run without harmonic restraints. Production runs were performed with an increased time step of 2 fs under the same ensemble. The electrostatic interactions were calculated using the Particle Mesh Ewald (PME) method and all bonds to hydrogen atoms were constrained using the SHAKE algorithm. For each P-gp nucleotide system, 120 replicas of 10 ns production runs were performed starting from randomly assigned initial velocities, where the RMSDs of fourteen segments, including twelve TM helices and two NBDs, were computed with respect to the first frame. Among the 120 replicas, domain-specific RMSD outlier detection (Q3 + 1.5×IQR) identified replicas exploring under-sampled substates, which were extended for 90 ns. Total production run from all P-gp conformers and nucleotide states was ∼110 μs (Table 1,2).

### 3. Analysis of Flexible Linker Interaction

To conduct principal component (PC) analysis, all replica trajectories were featurized with pyEMMA (20) using the closest heavy atom distances between the residues 159-189, 630-700, 801-831 to include intracellular portions of TM3,9,10 and the flexible linker. The percentage contributions of the top five PCs were 77.15, 6.27, 3.29, 1.73, 1.35, respectively. The first two PCs were used to generate 2D histograms, which were constructed as heatmaps using kernel density estimation. Residue-wise secondary structure analysis was conducted using the DSSP algorithm (67) and CPPTRAJ (68).

### 4. Binding-Free Energy Calculations

Binding free energies for nucleotides (ATP/ADP) were estimated by the molecular-mechanics/Poisson-Boltzmann surface-area (MM/PBSA) protocol in 0.15 M salt concentration (69). Prior to the energy decomposition, membrane lipids, ions and water in the original system were stripped so that only protein atoms were considered as receptor. Fifty uncorrelated snapshots were extracted from each 100 ns trajectory.

### 5. CAVER Tunnel Analysis

The CAVER 3.0 software (21) was used to identify the possible ligand access tunnels in the simulation trajectories with frames saved at intervals of 200 ps. Initial 30 percent of the frames of each of the trajectories were stripped to minimize starting conformation bias. All frames of the trajectories belonging to the unique P-gp conformer and nucleotide state were combined and aligned to an initial frame. For all CAVER analyses, tunnel calculation was performed excluding the residues 1-44, 370-710, 1013-1280 to only include the TMD region. The starting points were defined as the benzene carbons in F336 and F983 to represent the canonical ligand binding site. CAVER tunnel analysis was run using a probe radius of 2.5 Å, shell radius of 6.0 Å, and shell-depth of 4.0 Å.

## Author contributions

SP, HF and JW jointly conceived and supervised the study. SBH designed the study, performed calculations, analyzed the data, and wrote the manuscript. SBH, SP, HF and JW jointly edited the manuscript.

## Data availability

The input files and last frames of all MD simulation replicas and CAVER analysis are deposited and available on Zenodo at https://doi.org/10.5281/zenodo.15479648.

## Acknowledgements

The authors gratefully acknowledge the support of the A*STAR Research Attachment Programme (ARAP) scholarship and the provision of computing resources by the N8 CIR through Bede national tier-2 supercomputer facility in the United Kingdom, and by the Computational Shared Facility (CSF3) at the University of Manchester, and by the high-performance computing facility, National Supercomputing Centre (NSCC) Singapore. The authors thank Dave Love for assistance with technical aspects of the computations at Bede supercomputer and Dr Alessandro Barbieri for helpful discussions.

## References

1. Thomas C, Tampé R. Structural and Mechanistic Principles of ABC Transporters. Annu Rev Biochem. 2020;89(1):605–36.

2. Rees DC, Johnson E, Lewinson O. ABC Transporters: The Power to Change. Nat Rev Mol Cell Biol. 2009;10:218–27.

3. Schneider E. ATP-Binding Cassette Transport Systems: Functional and Structural Aspects of the ATP-Hydrolyzing Subunits. FEMS Microbiol Rev. 1998;22(1):1–20.

4. Ford RC, Beis K. Learning the ABCs One at a Time: Structure and Mechanism of ABC Transporters. Biochem Soc Trans. 2019;47:23–36.

5. Juliano RL, Ling V. A Surface Glycoprotein Modulating Drug Permeability in Chinese Hamster Ovary Cell Mutants. Biochim Biophys Acta. 1976;455:152–62.

6. Tian Y, Lei Y, Wang Y, Lai J, Wang J, Xia F. Mechanism of Multidrug Resistance to Chemotherapy Mediated by P-Glycoprotein. Int J Oncol. 2023;63(5):1019–29.

7. Chen Z, Shi T, Zhang L, Zhu P, Deng M, Huang C, et al. Mammalian Drug Efflux Transporters of the ATP Binding Cassette Family in Multidrug Resistance: A Review of the Past Decade. Cancer Lett. 2016;370(1):153–64.

8. Ambudkar SV, Kimchi-Sarfaty C, Sauna ZE, Gottesman MM. P-Glycoprotein: From Genomics to Mechanism. Oncogene. 2003;22(47):7468–85.

9. Szewczyk P, Tao H, McGrath AP, Villaluz M, Rees SD, Lee SC, et al. Snapshots of Ligand Entry, Malleable Binding and Induced Helical Movement in P-Glycoprotein. Acta Crystallogr Biol Crystallogr. 2015;71(3):732–41.

10. Li J, Jaimes KF, Aller SG. Refined Structures of Mouse P-Glycoprotein. Protein Sci. 2014;23(1):34–46.

11. Culbertson AT, Liao M. Cryo-EM of Human P-Glycoprotein Reveals an Intermediate Occluded Conformation During Active Drug Transport. Nat Commun. 2025;16:3619.

12. Jones PM, O’Mara ML, George AM. ABC Transporters: A Riddle Wrapped in a Mystery Inside an Enigma. Trends Biochem Sci. 2009;34:520–31.

13. Beis K. Structural Basis for the Mechanism of ABC Transporters. Biochem Soc Trans. 2015;43(5):889–93.

14. Moeller A, Lee SC, Tao H, Speir JA, Chang G, Urbatsch IL, et al. Distinct Conformational Spectrum of Homologous Multidrug ABC Transporters. Structure. 2015;23(3):450–60.

15. Frank GA, Shukla S, Rao P, Borgnia MJ, Bartesaghi A, Merk A, et al. Cryo-EM Analysis of the Conformational Landscape of Human P-Glycoprotein (ABCB1) During Its Catalytic Cycle. Mol Pharmacol. 2016;90(1):35–41.

16. Garrigues A, Escargueil AE, Orlowski S. The Multidrug Transporter P-Glycoprotein Actively Mediates Cholesterol Redistribution in the Cell Membrane. Proc Natl Acad Sci USA. 2002;99(16):10347–52.

17. Verhalen B, Dastvan R, Thangapandian S, et al. Energy Transduction and Alternating Access of the Mammalian ABC Transporter P-Glycoprotein. Nature. 2017;543(7647):738–41.

18. Dawson RJP, Locher KP. Structure of a Bacterial Multidrug ABC Transporter. Nature. 2006;443(7108):180–5.

19. Kim Y, Chen J. Molecular Structure of Human P-Glycoprotein in the ATP-Bound, Outward-Facing Conformation. Science. 2018;359(6378):915–9.

20. Scherer MK, Trendelkamp-Schroer B, Paul F, Pérez-Hernández G, Hoffmann M, Plattner N, et al. PyEMMA 2: A Software Package for Estimation, Validation, and Analysis of Markov Models. J Chem Theory Comput. 2015;11(11):5525–42.

21. Chovancova E, Pavelka A, Benes P, Strnad O, Brezovsky J, Kozlikova B, et al. CAVER 3.0: A Tool for the Analysis of Transport Pathways in Dynamic Protein Structures. PLoS Comput Biol. 2012;8(10):e1002708.

22. Jumper J, Evans R, Pritzel A, Green T, Figurnov M, Ronneberger O, et al. Highly Accurate Protein Structure Prediction with AlphaFold. Nature. 2021;596(7873):583–9.

23. Wan S, Bhati AP, Wade AD, Coveney PV. Ensemble-Based Approaches Ensure Reliability and Reproducibility. J Chem Theory Comput. 2023;19(22):6959–63.

24. Knapp B, Ospina L, Deane CM. Avoiding False Positive Conclusions in Molecular Simulation: The Importance of Replicas. J Chem Theory Comput. 2018;14(12):6127–38.

25. Tia H, Jiang X, Xiao S, Force LF, Larson EC, Tao P. LAST: Latent Space Assisted Adaptive Sampling for Protein Trajectories. J Chem Inf Model. 2022;63(1):67–75.

26. Condic-Jurkic K, Subramanian N, Mark AE, O’Mara ML. The Reliability of Molecular Dynamics Simulations of the Multidrug Transporter P-Glycoprotein in a Membrane Environment. PLoS One. 2018;13(1):e0191882.

27. Wang L, O’Mara ML. Effect of the Force Field on Molecular Dynamics Simulations of the Multidrug Efflux Protein P-Glycoprotein. J Chem Theory Comput. 2021;17(10):6491–508.

28. Stefan E, Hofmann S, Tampé R. A Single Power Stroke by ATP Binding Drives Substrate Translocation in a Heterodimeric ABC Transporter. Elife. 2020;9:e55943.

29. Dastvan R, Mishra S, Peskova YB, Nakamoto RK, Mchaourab HS. Mechanism of allosteric modulation of P-glycoprotein by transport substrates and inhibitors. Science. 2019 May 17;364(6441):689–92.

30. Bársony O, Szalóki G, Türk D, et al. A Single Active Catalytic Site Is Sufficient to Promote Transport in P-Glycoprotein. Sci Rep. 2016;6:24810.

31. Szöll\Hosi D, Chiba P, Szakacs G, Stockner T. Conversion of Chemical to Mechanical Energy by the Nucleotide Binding Domains of ABCB1. Sci Rep. 2020;10:5943.

32. Hrycyna CA, Ramachandra M, Germann UA, Cheng PW, Pastan I, Gottesman MM. Both ATP Sites of Human P-Glycoprotein Are Essential but Not Symmetric. Biochemistry. 1999;38(42):13887–99.

33. Ward A, Reyes CL, Yu J, Roth CB, Chang G. Flexibility in the ABC Transporter MsbA: Alternating Access with a Twist. Proc Natl Acad Sci USA. 2007;104(48):19005–10.

34. Oliveira AS, Baptista AM, Soares CM. Conformational Changes Induced by ATP Hydrolysis in an ABC Transporter: A Molecular Dynamics Study of the Sav1866 Exporter. Proteins. 2011;79(6):1977–90.

35. Pan L, Aller SG. Equilibrated Atomic Models of Outward-Facing P-Glycoprotein and Effect of ATP Binding on Structural Dynamics. Sci Rep. 2015;5:7880.

36. Nosol K, Romane K, Irobalieva RN, Alam A, Kowal J, Fujita N, et al. Cryo-EM Structures Reveal Distinct Mechanisms of Inhibition of the Human Multidrug Transporter ABCB1. Proc Natl Acad Sci USA. 2020;117(42):26245–53.

37. Choudhury HG, Tong Z, Mathavan I, et al. Structure of an Antibacterial Peptide ABC Transporter in a Novel Outward Occluded State. Proc Natl Acad Sci USA. 2014;111(25):9145–50.

38. Best RB, de Sancho D, Mittal J. Residue-Specific Alpha-Helix Propensities from Molecular Simulation. Biophys J. 2012;102(6):1462–7.

39. Germann UA. Molecular Analysis of the Multidrug Transporter. Cytotechnology. 1993;12(1–3):33–62.

40. Hrycyna CA, Ramachandra M, Ambudkar SV, et al. Structural Flexibility of the Linker Region of Human P-Glycoprotein Permits ATP Hydrolysis and Drug Transport. Biochemistry. 1998;37(39):13660–73.

41. Germann UA, Chambers TC, Ambudkar SV, et al. Characterization of Phosphorylation-Defective Mutants of Human P-Glycoprotein Expressed in Mammalian Cells. J Biol Chem. 1996;271(3):1708–16.

42. Chambers TC, Pohl J, Raynor RL, Kuo JF. Identification of Specific Sites in Human P-Glycoprotein Phosphorylated by Protein Kinase C. J Biol Chem. 1993;268(7):4592–5.

43. Ford RC, Marshall-Sabey D, Schuetz J. Linker Domains: Why ABC Transporters “Live in Fragments No Longer”. Trends Biochem Sci. 2020;45(2):137–48.

44. Gottesman MM, Hrycyna CA, Schoenlein PV, et al. Genetic Analysis of the Multidrug Transporter. Annu Rev Genet. 1995;29:607–49.

45. Bonito CA, Ferreira RJ, Ferreira MJU, et al. Theoretical Insights on Helix Repacking as the Origin of P-Glycoprotein Promiscuity. Sci Rep. 2020;10:66587.

46. Ferreira RJ, Ferreira MJU, dos Santos DJVA. Insights on P-Glycoprotein’s Efflux Mechanism Obtained by Molecular Dynamics Simulations. J Chem Theory Comput. 2012;8(6):1853–64.

47. Alam A, Küng R, Kowal J, et al. Structure of the Zosuquidar-Bound Human P-Glycoprotein Reveals a Dual-Mode Inhibitor. Nature. 2019;573(7773):223–6.

48. Kurre D, Dang PX, Le LTM, et al. Structural Insights into Binding-Site Access and Ligand Recognition by Human ABCB1. EMBO J. 2025;44:991–1006.

49. Hamaguchi-Suzuki N, Adachi N, Moriya T, et al. Cryo-EM Structure of P-Glycoprotein Bound to Triple Elacridar Inhibitor Molecules. Biochem Biophys Res Commun. 2024;709:149855.

50. Higgins C, Linton K. The ATP Switch Model for ABC Transporters. Nat Struct Mol Biol. 2004;11:918–26.

51. Becker JP, Van Bambeke F, Tulkens PM, Prévost M. Dynamics and Structural Changes Induced by ATP Binding in SAV1866, a Bacterial ABC Exporter. J Phys Chem B. 2010;114(48):15948–57.

52. Xu Y, Seelig A, Bernèche S. Unidirectional Transport Mechanism in an ATP Dependent Exporter. ACS Cent Sci. 2017;3(3):250–8.

53. Lusvarghi S, Durell SR, Ambudkar SV. Does the ATP-Bound EQ Mutant Reflect the Pre- or Post-ATP Hydrolysis State in the Catalytic Cycle of Human P-Glycoprotein (ABCB1)? FEBS Lett. 2021;595(6):750–62.

54. Lacabanne D, Wiegand T, De Cesare M, Orelle C, Ernst M, Jault J, et al. Solid-State NMR Reveals Asymmetric ATP Hydrolysis in the Multidrug ABC Transporter BmrA. J Am Chem Soc. 2022;144(27):12431–42.

55. Vigano C, Julien M, Carrier I, Gros P, Ruysschaert JM. Structural and Functional Asymmetry of the Nucleotide-Binding Domains of P-Glycoprotein Investigated by Attenuated Total Reflection FTIR Spectroscopy. J Biol Chem. 2002;277(7):5008–16.

56. Eswar N, Webb B, Marti-Renom MA, Madhusudhan MS, Eramian D, Shen MY, et al. Comparative Protein Structure Modeling Using MODELLER. In: Curr Protoc Protein Sci. 2007. p. Unit\ 2.9.

57. Wu EL, Cheng X, Jo S, Rui H, Song KC, Dávila-Contreras EM, et al. CHARMM-GUI Membrane Builder Toward Realistic Biological Membrane Simulations. J Comput Chem. 2014;35:1997–2004.

58. Lomize MA, Pogozheva ID, Joo H, Mosberg HI, Lomize AL. OPM Database and PPM Web Server: Resources for Positioning of Proteins in Membranes. Nucleic Acids Res. 2012;40:D370–6.

59. Shintre CA, Pike ACW, Li Q, Kim JI, Barr AJ, Goubin S, et al. Structures of ABCB10, a Human ATP-Binding Cassette Transporter in Apo- and Nucleotide-Bound States. Proc Natl Acad Sci USA. 2013;110(24):9710–5.

60. Maier JA, Martinez C, Kasavajhala K, Wickstrom L, Hauser KE, Simmerling C. ff14SB: Improving the Accuracy of Protein Side Chain and Backbone Parameters from ff99SB. J Chem Theory Comput. 2015;11(8):3696–713.

61. Dickson CJ, Madej BD, Walker RC. Lipid14: The AMBER Lipid Force Field. J Chem Theory Comput. 2014;10:865–79.

62. Meagher KL, Redman LT, Carlson HA. Development of polyphosphate parameters for use with the AMBER force field. J Comput Chem. 2003 Jul 15;24(9):1016–25.

63. Allnér O, Nilsson L, Villa A. Magnesium Ion-Water Coordination and Exchange in Biomolecular Simulations. J Chem Theory Comput. 2012;8(4):1493–502.

64. Case DA, Belfon K, Ben-Shalom IY, et al. AMBER 2020. University of California, San Francisco; 2020.

65. Han SB, Teuffel J, Mukherjee G, Wade RC. Multiresolution molecular dynamics simulations reveal the interplay between conformational variability and functional interactions in membrane-bound cytochrome P450 2B4. Protein Sci. 2024 Oct;33(10):e5165.

66. Abraham MJ, Murtola T, Schulz R, Päll S, Smith JC, Hess B, et al. GROMACS: High Performance Molecular Simulations Through Multi-Level Parallelism from Laptops to Supercomputers. SoftwareX. 2015;1–2:19–25.

67. Kabsch W, Sander C. Dictionary of Protein Secondary Structure: Pattern Recognition of Hydrogen-Bonded and Geometrical Features. Biopolymers. 1983;22(12):2577–637.

68. Roe DR, Cheatham TE. PTRAJ and CPPTRAJ: Software for Processing and Analysis of Molecular Dynamics Trajectory Data. J Chem Theory Comput. 2013;9(7):3084– 95.

69. Srinivasan J, Cheatham TE, Cieplak P, Kollman PA, Case DA. Continuum Solvent Studies of the Stability of DNA, RNA, and Phosphoramidate−DNA Helices. J Am Chem Soc. 1998 Sep 1;120(37):9401–9.

